# A Library of Nucleotide Analogues Terminate RNA Synthesis Catalyzed by Polymerases of Coronaviruses Causing SARS and COVID-19

**DOI:** 10.1101/2020.04.23.058776

**Authors:** Steffen Jockusch, Chuanjuan Tao, Xiaoxu Li, Thomas K. Anderson, Minchen Chien, Shiv Kumar, James J. Russo, Robert N. Kirchdoerfer, Jingyue Ju

**Affiliations:** Center for Genome T echnology and Biomolecular Engineering, Columbia University, New York, NY 10027; Departments of Chemistry, Columbia University, New York, NY 10027; Departments of Chemical Engineering, Columbia University, New York, NY 10027; Pharmacology, Columbia University, New York, NY 10027; Departments of Departments of Biochemistry, University of Wisconsin-Madison, Madison, WI 53706; Institute of Molecular Virology, University of Wisconsin-Madison, Madison, WI 53706

**Keywords:** COVID-19, SARS-CoV-2, RNA-dependent RNA polymerase, Nucleotide analogues, Exonuclease

## Abstract

SARS-CoV-2, a member of the coronavirus family, is responsible for the current COVID-19 worldwide pandemic. We previously demonstrated that five nucleotide analogues inhibit the SARS-CoV-2 RNA-dependent RNA polymerase (RdRp), including the active triphosphate forms of Sofosbuvir, Alovudine, Zidovudine, Tenofovir alafenamide and Emtricitabine. We report here the evaluation of a library of additional nucleoside triphosphate analogues with a variety of structural and chemical features as inhibitors of the RdRps of SARS-CoV and SARS-CoV-2. These features include modifications on the sugar (2’ or 3’ modifications, carbocyclic, acyclic, or dideoxynucleotides) or on the base. The goal is to identify nucleotide analogues that not only terminate RNA synthesis catalyzed by these coronavirus RdRps, but also have the potential to resist the viruses’ exonuclease activity. We examined these nucleotide analogues with regard to their ability to be incorporated by the RdRps in the polymerase reaction and then prevent further incorporation. While all 11 molecules tested displayed incorporation, 6 exhibited immediate termination of the polymerase reaction (Carbovir triphosphate, Ganciclovir triphosphate, Stavudine triphosphate, Entecavir triphosphate, 3’-O-methyl UTP and Biotin-16-dUTP), 2 showed delayed termination (Cidofovir diphosphate and 2’-O-methyl UTP), and 3 did not terminate the polymerase reaction (2’-fluoro-dUTP, 2’-amino-dUTP and Desthiobiotin-16-UTP). The coronavirus genomes encode an exonuclease that apparently requires a 2’ -OH group to excise mismatched bases at the 3’-terminus. In this study, all of the nucleoside triphosphate analogues we evaluated form Watson-Crick-like base pairs. All the nucleotide analogues which demonstrated termination either lack a 2’-OH, have a blocked 2’-OH, or show delayed termination. These nucleotides may thus have the potential to resist exonuclease activity, a property that we will investigate in the future. Furthermore, prodrugs of five of these nucleotide analogues (Brincidofovir/Cidofovir, Abacavir, Valganciclovir/Ganciclovir, Stavudine and Entecavir) are FDA approved for other viral infections, and their safety profile is well known. Thus, they can be evaluated rapidly as potential therapies for COVID-19.

## 1. Introduction

The COVID-19 pandemic, caused by SARS-CoV-2, continues to have a devastating global impact. SARS-CoV-2 is a member of the Orthocoronavirinae subfamily (Zhu et al. 2020). Coronaviruses, HCV and the flaviviruses are all positive-sense single-strand RNA viruses that replicate their genomes using an RNA-dependent RNA polymerase (RdRp) catalyzed reaction (Zumla et al. 2016; Dustin et al. 2016).

Currently, there are no FDA approved antiviral drugs for the treatment of human coronavirus infections, including COVID-19. The RdRp of coronaviruses is a well established drug target; the active site of the RdRp is highly conserved among positive-sense RNA viruses (te Velthuis 2014). These RdRps have low fidelity (Selisko et al. 2018), allowing them to recognize a variety of modified nucleotide analogues as substrates. Such nucleotide analogues may inhibit further RNA-polymerase catalyzed RNA replication making them important candidate anti-viral agents (McKenna et al. 1989; Öberg 2006; Eltahla et al. 2015; De Clercq and Li 2016). RdRps in SARS-CoV and SARS-CoV-2 have nearly identical sequences (Elfiky 2020); recently, the SARS-CoV-2 RdRp was cloned (Chien et al. 2020) and the RNA polymerase complex structure was determined (Gao et al. 2020), which will help guide the design and study of RdRp inhibitors.

Remdesivir, a phosphoramidate prodrug containing a 1’-cyano modification on the sugar, is converted in cells into an adenosine triphosphate analogue, which has been shown to be an inhibitor of the RdRp of SARS-CoV and SARS-CoV-2 (Gordon et al. 2020a; 2020b). It is currently in clinical trials in several countries as a therapeutic for COVID-19 infections. Remdesivir triphosphate has been shown to be incorporated with higher efficiency than ATP by coronavirus RdRps; it shows delayed termination which could help to overcome excision by the viral exonuclease (Gordon et al. 2020a; 2020b). β-D-N^4^-hydroxycytidine is another prodrug targeting the polymerase and has been shown to have broad spectrum activity against coronaviruses, even in the presence of intact proofreading functions (Agostini et al. 2019, Sheahan et al. 2020).

### 1.1. Selection of candidate nucleoside triphosphates for Coronavirus polymerase testing

We previously demonstrated that five nucleotide analogues inhibit the SARS-CoV-2 RNA-dependent RNA polymerase (RdRp), including the active triphosphate forms of Sofosbuvir, Alovudine, Zidovudine, Tenofovir alafenamide and Emtricitabine (Ju et al. 2020; Chien et al. 2020; Jockusch et al. 2020). Emtricitabine and Tenofovir alafenamide are used in FDA approved combination regimens for treatment of HIV/AIDS and hepatitis B virus (HBV) infections as well as pre-exposure prophylaxis (PrEP) to prevent HIV infections (Anderson et al. 2011), and may be evaluated for PrEP for COVID-19 (Jockusch et al. 2020).

The fact that all of the previous five nucleotide analogues exhibited inhibition of the coronavirus polymerases indicates that the SARS-CoV-2 RdRp can accept a variety of nucleotide analogues as substrates. In this work, we evaluate additional nucleotide analogues with a larger variety of modifications to identify those with more efficient termination, as well as the ability to resist the virus’s proofreading function. These additional nucleotide analogues were selected based on one or more of the following criteria. First, they have structural and chemical properties such as (a) similarity in size and structure to natural nucleotides, including the ability to fit within the active site of the polymerase, (b) presence of modifications at the 3’-OH position or absence of a 3’-OH group resulting in obligate termination of the polymerase reaction; or (c) modifications at the 2’ or other positions on the sugar or base that can delay termination. The above criteria will provide structural and chemical features that we can explore to evade exonuclease activity (Minskaia et al. 2006). Second, if they have previously been shown to inhibit the polymerases of other viruses, they may also have the potential to inhibit the SARS-CoV-2 polymerase based on our previous work. Third, ideally, the inhibitors should display high selectivity for viral polymerases relative to cellular DNA or RNA polymerases. And fourth, there is an advantage in considering nucleotide analogues that are the active triphosphate forms of FDA-approved drugs, as these drugs are known to have acceptable levels of toxicity and are more likely to be tolerated by patients with coronavirus infections, including COVID-19.

Using the above selection criteria, here we examine 11 nucleotide analogues with sugar or base modifications (structures shown in Fig. 1) for their ability to inhibit the SARS-CoV-2 or SARS-CoV RdRps: Ganciclovir 5’-triphosphate, Carbovir 5’-triphosphate, Cidofovir diphosphate, Stavudine 5’-triphosphate, Entecavir 5’-triphosphate, 2’-O-methyluridine-5’-triphosphate, 3’-O-methyluridine-5’-triphosphate, 2’-fluoro-2’ -deoxyuridine-5’ -triphosphate, desthiobiotin-16-aminoallyl-2’ -uridine-5’ - triphosphate, biotin-16-aminoallyl-2’-deoxyuridine-5’-triphosphate, and 2’-aminouridine-5’-triphosphate. The nucleoside and prodrug forms for the FDA approved drugs are shown in Fig. 2, and nucleoside and potential prodrug forms for three of the others are shown in Fig. S1.

**Fig. 1.**
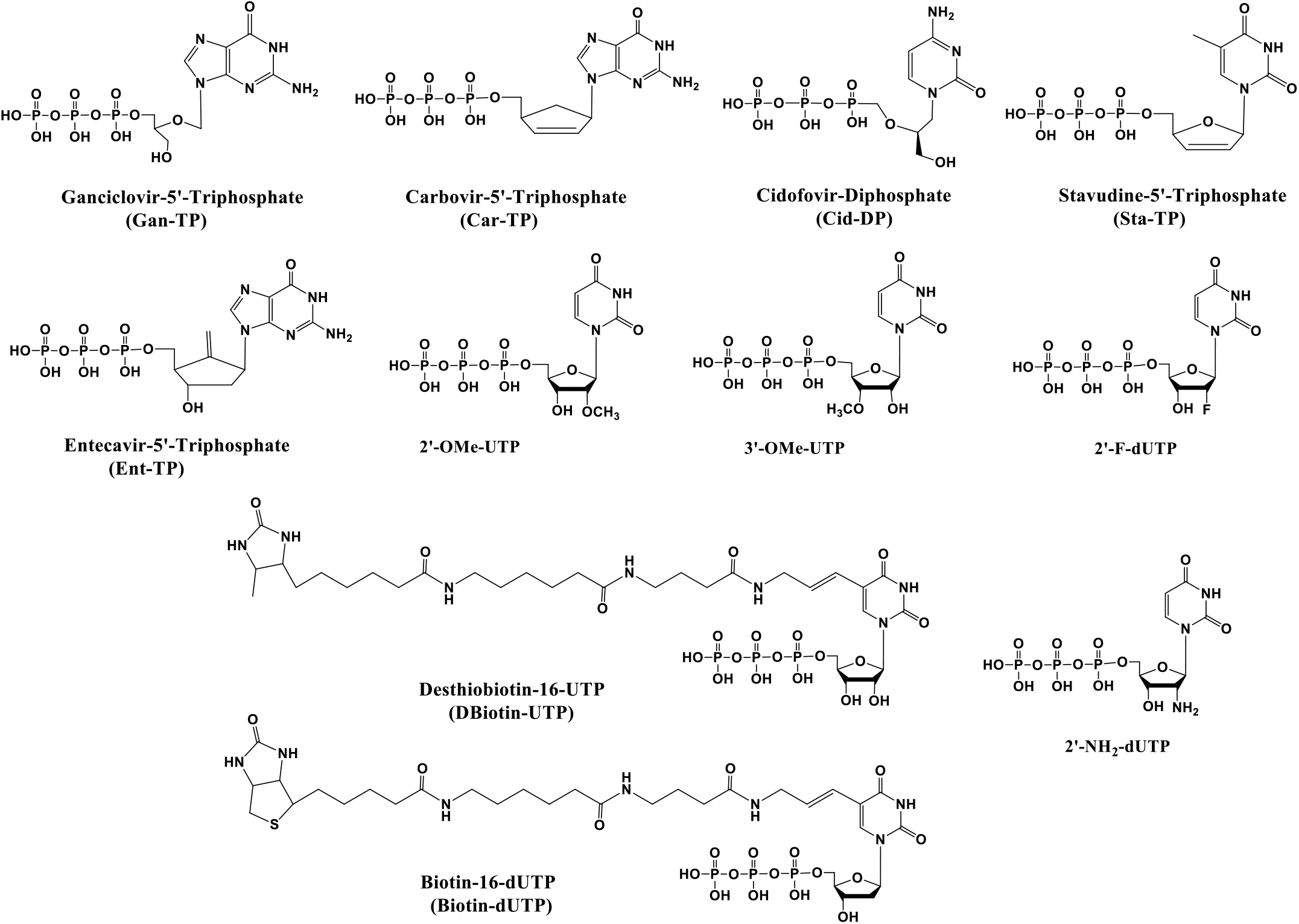
Chemical structures of nucleoside triphosphate analogues used in this study.

**Fig. 2.**
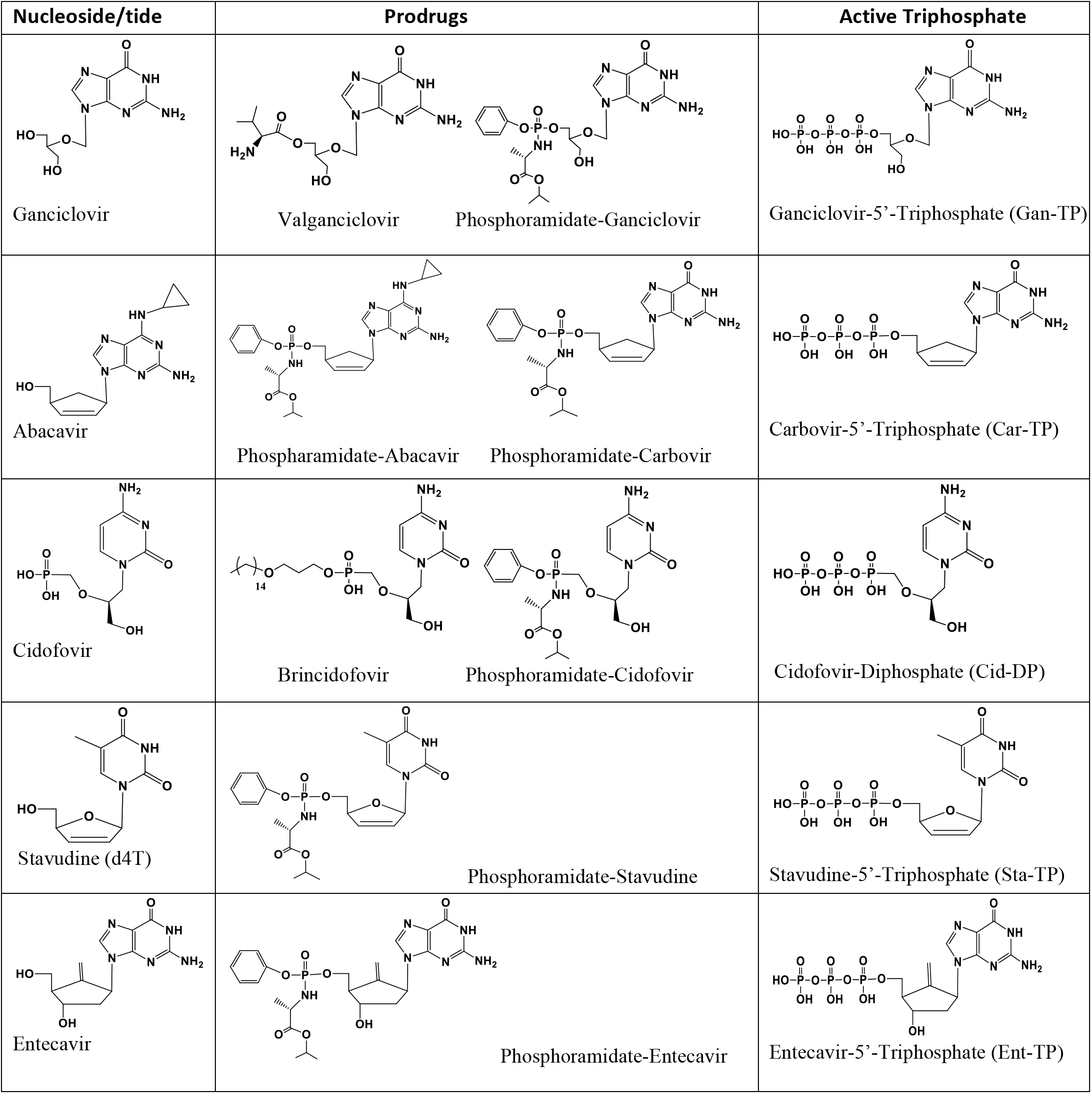
Structures of viral nucleoside/nucleotide inhibitors, example prodrugs and active triphosphate forms. The compounds Ganciclovir, Abacavir, Cidofovir, Stavudine and Entecavir (left), example prodrug forms (middle) and their active triphosphate forms (right).

Some of the uridine analogues listed above have been investigated as inhibitors of viral polymerases (Kumaki et al. 2011; Eyer et al. 2019; Arup et al. 1992; Lauridsen et al. 2012). The 2’-O-methyluridine triphosphate is of particular interest, since it has been demonstrated that 2’-O-methyl nucleotides are resistant to removal by the 3’-exonuclease found in coronaviruses (Minskaia et al. 2006). Next, we briefly describe 5 nucleotide analogues whose prodrug forms have been FDA approved for other virus infections.

Ganciclovir triphosphate (Gan-TP) is an acyclic guanosine nucleotide (Fig. 1). The parent nucleoside Ganciclovir (Cytovene, Fig. 2) is used to treat AIDS-related cytomegalovirus (CMV) infections. The drug can inhibit herpesviruses including herpesvirus 1 and 2, varicella zoster virus and CMV *in vitro* and *in vivo.* The valyl ester prodrug Valganciclovir (Fig. 2) can be given orally. After cleavage of the valyl ester, Ganciclovir is converted to Ganciclovir triphosphate by viral and cellular enzymes (Matthews and Boehme 1988; Akyürek et al. 2001).

Carbovir triphosphate (Car-TP) is a carbocyclic guanosine didehydro-dideoxynucleotide (Fig. 1). The parent prodrug is Abacavir (Ziagen, Fig. 2), which is an FDA approved nucleoside reverse transcriptase inhibitor for HIV/AIDS treatment (Faletto et al. 1997; Ray et al. 2002). It is taken orally and is well tolerated.

Cidofovir diphosphate (Cid-DP) is an acyclic cytidine nucleotide (Fig. 1). Its prodrug form Cidofovir (Vistide, Fig. 2) has been FDA approved for the intravenous treatment of AIDS-related CMV retinitis, and has been used off-label for a variety of DNA virus infections (De Clercq 2002; Lanier et al. 2010). The parent prodrug Brincidofovir (BDV, Fig. 2) is an oral antiviral drug candidate for treating smallpox infections. It has a lipid moiety masking the phosphate group, is taken up well by cells, has low toxicity, and is active against a wide range of DNA viruses in animals, including poxviruses, adenoviruses, herpesviruses, and CMV (Trost et al. 2015; Cundy et al. 1999). Interestingly, it has been shown using vaccinia virus DNA polymerase, that though Cid-DP is incorporated into DNA in the polymerase reaction, the process is relatively inefficient, and termination of synthesis occurs after extension by an additional nucleotide, a delayed termination similar to that shown for Remdesivir for RdRp; in the penultimate position, the incorporated Cidofovir is not removed by 3’-exonuclease activity and resistance is not common for Cidofovir (Magee et al. 2005).

Stavudine triphosphate (Fig. 1), a thymidine analogue, is the active triphosphate form of Stavudine (d4T, Zerit, Fig. 2), an antiviral used for the prevention and treatment of HIV/AIDS (Ho and Hitchcock 1989). It preferentially inhibits the HIV reverse transcriptase (RT) (Huang et al. 1992). The lack of a 3’-OH group makes it an obligate inhibitor.

Entecavir triphosphate (Ent-TP, Fig. 1), the active triphosphate form of the orally available drug Entecavir (Baraclude, Fig. 2), is a guanosine nucleotide inhibitor of the HBV RT (Matthews 2006, Rivkina and Rybalow 2002). It shows little if any inhibition of nuclear and mitochondrial DNA polymerases (Mazzucco et al. 2008), and has generally been shown to have low toxicity. It has been shown that Entecavir triphosphate is a delayed chain terminator with the HIV-1 reverse transcriptase, making it resistant to excision by exonucleases (Tchesnokov et al. 2008).

We reasoned that once the above nucleotide analogues are incorporated into an RNA primer in the polymerase reaction, the fact that they either lack a normal sugar ring configuration or lack the 2’- and/or 3’-OH groups, would make it unlikely for them to be recognized by 3’-exonucleases involved in SARS-CoV-2 proofreading processes, decreasing the likelihood of developing resistance to inhibition by the drug.

### 1.2. Coronaviruses have a proofreading exonuclease activity that must be overcome to develop effective RdRp nucleotide inhibitors

In contrast to many other RNA viruses, the SARS-CoV and SARS-CoV-2 coronaviruses have very large genomes that encode a 3’-5’ exonuclease (nsp14) involved in proofreading (Shannon et al. 2020; Ma et al. 2015), the activity of which is enhanced by the cofactor nsp10 (Bouvet et al. 2012). This proofreading function increases replication fidelity by removing mismatched nucleotides (Ferron et al. 2018). Mutations in nsp14 lead to reduced replication fidelity of the viral genome (Eckerle et al. 2010). Interestingly, while the nsp14/nsp10 complex efficiently excises single mismatched nucleotides at the 3’ end of the RNA chain, it is not able to remove longer stretches of unpaired nucleotides or 3’ modified RNA (Bouvet et al. 2012). In order for the nucleotide analogues to be successful inhibitors of these viruses, they need to overcome this proofreading function. The coronavirus exonuclease activity typically requires the presence of a 2’-OH group at the 3’ end of the growing RNA strand (Minskaia et al. 2006). However, if there is delayed termination and the offending nucleotide analogue is no longer at the 3’ end, they will also not be removed by the exonuclease (Bouvet et al. 2012; Gordon et al. 2020a; 2020b). Nearly all of the nucleotide analogues we selected lack the 2’-OH group, have modifications that block the 2’-OH group on the sugar, or are acyclic nucleotide derivatives; such nucleotides will not likely be substrates for viral exonucleases.

## 2. Materials and Methods

### 2.1. Materials

Nucleoside triphosphates and nucleoside triphosphate analogues were purchased from TriLink BioTechnologies (Biotin-16-dUTP, Desthiobiotin-16-UTP, 2’-OMe-UTP, 3’-OMe-UTP, 2’-F-dUTP, 2’-NH_2_-dUTP, Cidofovir-DP, Ganciclovir-TP, dUTP, CTP, ATP and UTP), Santa Cruz Biotechnology (Stavudine-TP, Carbovir-TP), or Moravek, Inc. (Entecavir-TP). Oligonucleotides were purchased from Integrated DNA Technologies, Inc. (IDT).

### 2.2. Extension reactions with SARS-CoV-2 RNA-dependent RNA polymerase

The primer and template (sequences shown in Figs. 2–4, S2–S8) were annealed by heating to 70°C for 10 min and cooling to room temperature in 1x reaction buffer. The RNA polymerase mixture consisting of 6 *μ*M nsp12 and 18 *μ*M each of cofactors nsp7 and nsp8 (Chien et al. 2020) was incubated for 15 min at room temperature in a 1:3:3 ratio in 1x reaction buffer. Then 5 μl of the annealed template primer solution containing 2 *μ*M template and 1.7 *μ*M primer in 1x reaction buffer was added to 10 μl of the RNA polymerase mixture and incubated for an additional 10 min at room temperature. Finally 5 μl of a solution containing 2 mM Biotin-dUTP (a), 2 mM DBiotin-UTP (b), 2 mM 2’-OMe-UTP (c), 2 mM Sta-TP (d), 2 mM Cid-DP + 2 mM UTP + 2 mM ATP (e), 2 mM Car-TP + 2 mM UTP + 2 mM ATP + 2 mM CTP (f), 2 mM Gan-TP + 2 mM UTP + 2 mM ATP + 2 mM CTP (g), 2 mM Ent-TP + 2 mM UTP + 2 mM ATP + 2 mM CTP (h), 2 mM 2’-OMe-UTP + 2 mM dUTP (i), 1 mM UTP, 1 mM Biotin-dUTP and 1 mM dUTP (j), 1 mM 2’-F-dUTP, 1 mM 2’-OMe-UTP and 1 mM dUTP (k), or 1 mM 2’-NH_2_-dUTP, 1 mM 2’-OMe-UTP and 1 mM dUTP (l) in 1x reaction buffer was added, and incubation was carried out for 2 hrs at 30°C. The final concentrations of reagents in the 20 μl extension reactions were 3 *μ*M nsp12, 9 *μ*M nsp7, 9 *μ*M nsp8, 425 nM RNA primer, 500 nM RNA template, 500 *μ*M Biotin-dUTP (a), 500 *μ*M DBiotin-UTP (b), 500 *μ*M 2’-OMe-UTP (c), 500 *μ*M Sta-TP (d), 500 *μ*M Cid-DP, 500 *μ*M UTP and 500 *μ*M ATP (e), 500 *μ*M Car-TP, 500 *μ*M UTP, 500 *μ*M ATP and 500 *μ*M CTP (f), 500 *μ*M Gan-TP, 500 *μ*M UTP, 500 *μ*M ATP and 500 *μ*M CTP (g), 500 *μ*M Ent-TP + 500 *μ*M UTP + 500 *μ*M ATP + 500 *μ*M CTP (h), 500 *μ*M 2’-OMe-UTP + 500 *μ*M dUTP (i), 250 *μ*M UTP, 250 *μ*M Biotin-dUTP and 250 *μ*M dUTP (j), 250 *μ*M 2’-F-dUTP, 250 *μ*M 2’-OMe-UTP and 250 *μ*M dUTP (k), or 250 *μ*M 2’-NH_2_-dUTP, 250 *μ*M 2’-OMe-UTP and 250 *μ*M dUTP (l). The 1x reaction buffer contains the following reagents: 10 mM Tris-HCl pH 8, 10 mM KCl, 2 mM MgCl_2_ and 1 mM β-mercaptoethanol. Following desalting using an Oligo Clean & Concentrator (Zymo Research), the samples were subjected to MALDI-TOF-MS (Bruker ultrafleXtreme) analysis.

### 2.3. Extension reactions with SARS-CoV RNA-dependent RNA polymerase

The primer and template above were annealed by heating to 70°C for 10 min and cooling to room temperature in 1x reaction buffer (described above). The RNA polymerase mixture consisting of 6 *μ*M nsp12 and 18 *μ*M each of cofactors nsp7 and nsp8 (Kirchdoerfer and Ward 2019) was incubated for 15 min at room temperature in a 1:3:3 ratio in 1x reaction buffer. Then 5 μl of the annealed template primer solution containing 2 *μ*M template and 1.7 *μ*M primer in 1x reaction buffer was added to 10 μl of the RNA polymerase mixture and incubated for an additional 10 min at room temperature. Finally 5 μl of a solution containing 2 mM 2’-OMe-UTP (a), 2 mM 2’-F-dUTP (b), 2 mM 3’-OMe-UTP (c), 2 mM Cid-DP + 0.8 mM UTP + 0.8 mM ATP (d), 2 mM Car-TP + 0.8 mM UTP + 0.8 mM ATP + 0.8 mM CTP (e), or 2 mM Gan-TP + 0.8 mM UTP + 0.8 mM ATP + 0.8 mM CTP (f) in 1x reaction buffer was added, and incubation was carried out for 2 hrs at 30°C. The final concentrations of reagents in the 20 μl extension reactions were 3 *μ*M nsp12, 9 *μ*M nsp7, 9 *μ*M nsp8, 425 nM RNA primer, 500 nM RNA template, 500 *μ*M 2’-OMe-UTP (a), 500 *μ*M 2’-F-dUTP (b), 500 *μ*M 3’-OMe-UTP (c), 500 *μ*M Cid-DP, 200 *μ*M UTP and 200 *μ*M ATP (d), 500 *μ*M Car-TP, 200 *μ*M UTP, 200 *μ*M ATP and 200 *μ*M CTP (e), 500 *μ*M Gan-TP, 200 *μ*M UTP, 200 *μ*M ATP and 200 *μ*M CTP (f). Following desalting using an Oligo Clean & Concentrator, the samples were subjected to MALDI-TOF-MS analysis.

## 3. Results and Discussion

We tested the ability of the active triphosphate forms of the nucleotide analogues (structures shown in Fig. 1) to be incorporated by the RdRps of SARS-CoV or SARS-CoV-2. The prodrugs of these molecules are shown in Figs. 2 and S1. The phosphoramidate forms of prodrugs can be readily synthesized using the ProTide approach (Alanazi et al. 2019). The RdRp of these coronaviruses, referred to as nsp12, and its two protein cofactors, nsp7 and nsp8, have been shown to be required for the processive polymerase activity of nsp12 in SARS-CoV (Subissi et al. 2014; Kirchdoerfer and Ward 2019). These three components of each coronavirus polymerase complex were cloned and purified as described previously (Kirchdoerfer and Ward 2019; Chien et al. 2020). We then performed polymerase extension assays with 2’-O-methyluridine triphosphate (2’-OMe-UTP), 3’-O-methyluridine 5’-triphosphate (3’-OMe-UTP), 2’-fluoro-2’-deoxyuridine triphosphate (2’-F-dUTP), 2’-amino-2’-deoxyuridine triphosphate (2’-NH_2_-dUTP), biotin-16-dUTP (Biotin-UTP), desthiobiotin-16-UTP (DBiotin-UTP), Stavudine-TP (Sta-TP), Cidofovir diphosphate (Cid-DP) + UTP + ATP, Carbovir triphosphate (Car-TP) + UTP + ATP + CTP, Ganciclovir 5’-triphosphate (Gan-TP) + UTP + ATP + CTP, or Entecavir triphosphate (Ent-TP) + UTP + ATP + CTP, following the addition of a pre-annealed RNA template and primer to a pre-assembled mixture of the SARS-CoV and/or SARS-CoV-2 RdRp (nsp12) and the two cofactor proteins (nsp7 and nsp8). We also used combinations of nucleotide analogues in some cases to perform the polymerase reaction. The polymerase reaction products were analyzed by MALDI-TOF-MS. The sequences of the RNA template and primer used for the polymerase extension assay, which correspond to the 3’ end of the SARS-CoV-2 genome, are indicated at the top of Figs. 3–5 and S2–S8.

**Fig. 3.**
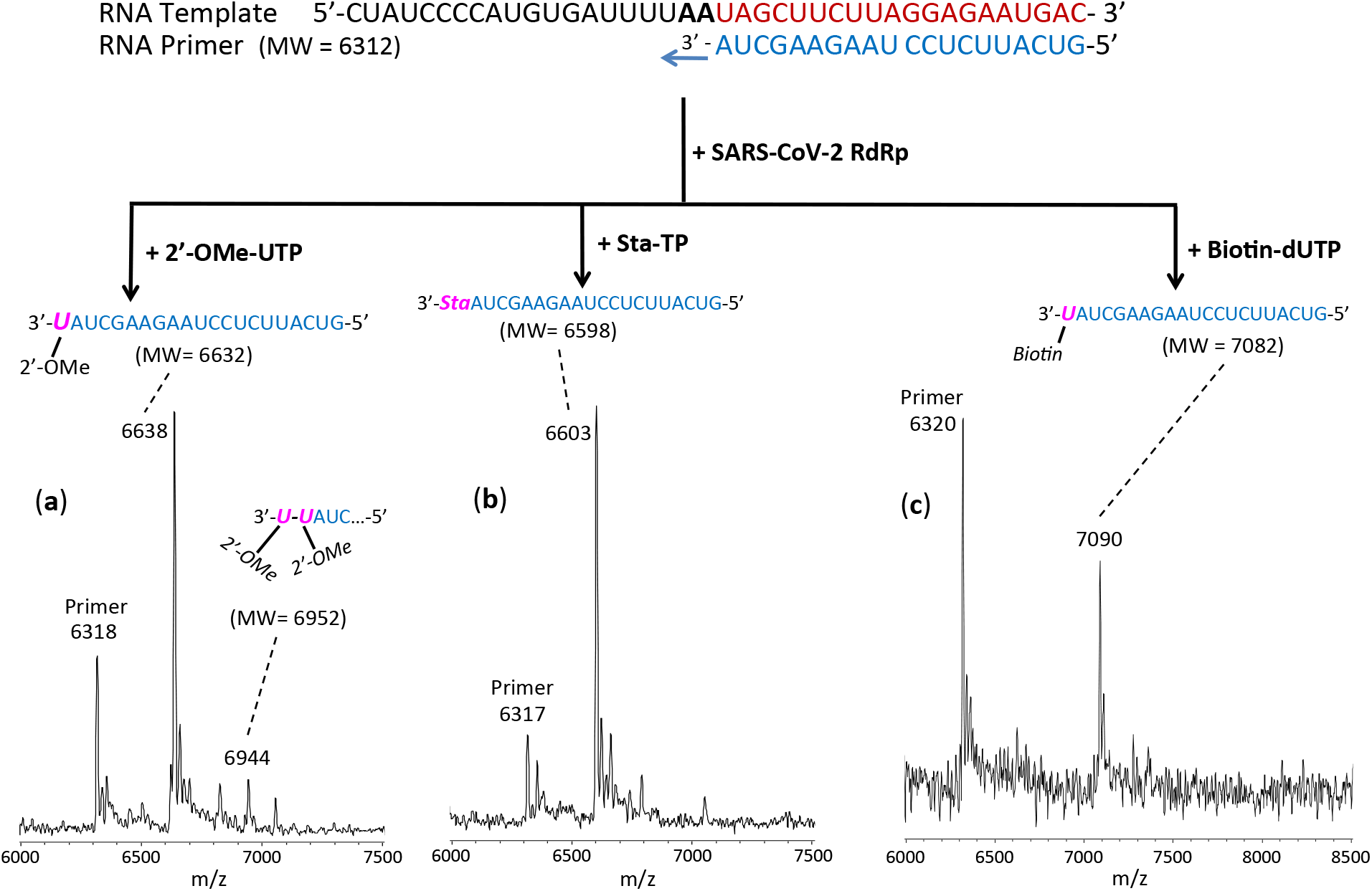
Incorporation of 2’-OMe-UTP, Sta-TP and Biotin-dUTP by SARS-CoV-2 RdRp to terminate the polymerase reaction. The sequences of the primer and template used for this extension reaction, which are at the 3’ end of the SARS-CoV-2 genome, are shown at the top of the figure. Polymerase extension reactions were performed by incubating 2’-OMe-UTP (a), Sta-TP (b) and Biotin-dUTP (c) with pre-assembled SARS-CoV-2 polymerase (nsp12, nsp7 and nsp8), the indicated RNA template and primer, and the appropriate reaction buffer, followed by detection of reaction products by MALDI-TOF MS. The detailed procedure is shown in the Methods section. The accuracy for m/z determination is ± 10 Da.

In the case of the UTP and TTP analogues, because there are two A’s in a row in the next available positions of the template for RNA polymerase extension downstream of the priming site, if they are indeed terminators of the polymerase reaction, the extension should stop after incorporating one nucleotide analogue. If they do not serve as terminators, two base extension by the UTP or TTP analogue will be observed. In the case of Cid-DP which is a CTP analogue, UTP and ATP must be provided to allow extension to the point where there is a G in the template strand. If the Cid-DP is then incorporated and acts as a terminator, extension will stop; otherwise, additional incorporation events may be observed. Similarly, for Carbovir-TP, Ganciclovir-TP, and Entecavir-TP, all of which are GTP analogues, UTP, ATP and CTP must be provided to allow extension to the point where there is a C in the template strand. If Car-TP, Gan-TP or Ent-TP is incorporated and acts as a terminator, extension will stop; otherwise additional incorporation events may be observed. Guided by polymerase extension results we obtained previously for the active triphosphate forms of Sofosbuvir, Alovudine, AZT, Tenofovir-DP and Emtricitabine-TP (Ju et al. 2020; Chien et al. 2020; Jockusch et al. 2020), various ratios of the nucleotides were chosen in the current work.

The results of the MALDI-TOF MS analysis of the primer extension reactions are shown in Figs. 3–5 and S2–S8. The observed peaks generally fit the nucleotide incorporation patterns described above; however, additional peaks assigned to intermediate stages of the extension reaction and in some cases extension beyond the incorporation of the nucleotide analogue was also observed. We describe the results for the SARS-CoV-2 polymerase catalyzed reaction in detail; similar results were obtained for the subset of nucleotide analogues tested with the SARS-CoV RdRp and are shown in the Supplementary Material.

The results for 2’-OMe-UTP, Sta-TP (which is a T analog) and Biotin-dUTP are presented in Fig. 3. In the case of extension with 2’-OMe-UTP, MS peaks representing incorporation by one 2’-OMe-UTP (6638 Da observed, 6632 Da expected) and to a lesser extent two 2’-OMe-UTPs (6944 Da observed, 6952 Da expected) were observed. Thus, 2’-OMe-UTP shows significant termination upon incorporation, indicating it can be a potential drug lead. 2’-O-methyl modification of RNA occurs naturally and therefore should have relatively low toxicity. In addition, ribose-2’-O-methylated RNA resists viral exonuclease activity (Minskaia et al. 2006). For Sta-TP, a single incorporation peak (6603 Da observed, 6598 Da expected) was seen, indicating that Sta-TP is very efficiently incorporated and achieves complete termination of the polymerase reaction. Since this molecule is a dideoxynucleotide without any hydroxyl groups on the sugar moiety, it may resist exonuclease activity. In the case of Biotin-dUTP, a single incorporation peak was evident (7090 Da observed, 7082 Da expected), suggesting that Biotin-dUTP is also a terminator of the polymerase reaction under these conditions. This indicates that the presence of a modification on the base along with the absence of a 2’-OH group in this nucleotide analogue leads to termination of the polymerase reaction catalyzed by the SARS-CoV-2 RdRp.

The result for the CTP analogue Cid-DP, which has an OH group, is presented in Fig. 4. Major peaks were observed indicating incorporation of a Cid-DP at the 8^th^ position from the initial priming site (8813 Da observed, 8807 Da expected) and a further 2 base extension by one ATP followed by one Cid-DP at the 10^th^ position (9404 Da observed, 9397 Da expected). There is no further extension beyond this position, indicative of delayed termination by Cid-DP. A small intermediate peak was also observed indicating extension by an ATP at the 9^th^ position from the initial priming site following the first Cid-DP incorporation (9142 Da observed, 9138 Da expected). A small partial UTP extension peak (6623 Da observed, 6618 Da expected) was also observed. An essentially identical result was obtained with the SARS-CoV polymerase (Fig. S2). Delayed termination for Cid-DP has been described for a vaccinia virus DNA polymerase (Magee et al. 2005). The investigational drug Remdesivir, currently being tested on COVID-19 patients, also displays delayed termination (Gordon et al. 2020a; 2020b); this is a major factor in its ability to resist the nsp14 3’-5’ exonuclease activity. Similar resistance to this exonuclease therefore will likely also occur with Cidofovir due to delayed termination of the polymerase reaction. Based on these results, Cidofovir and the oral prodrug Brincidofovir are thus potential leads for COVID-19 and SARS.

**Fig. 4.**
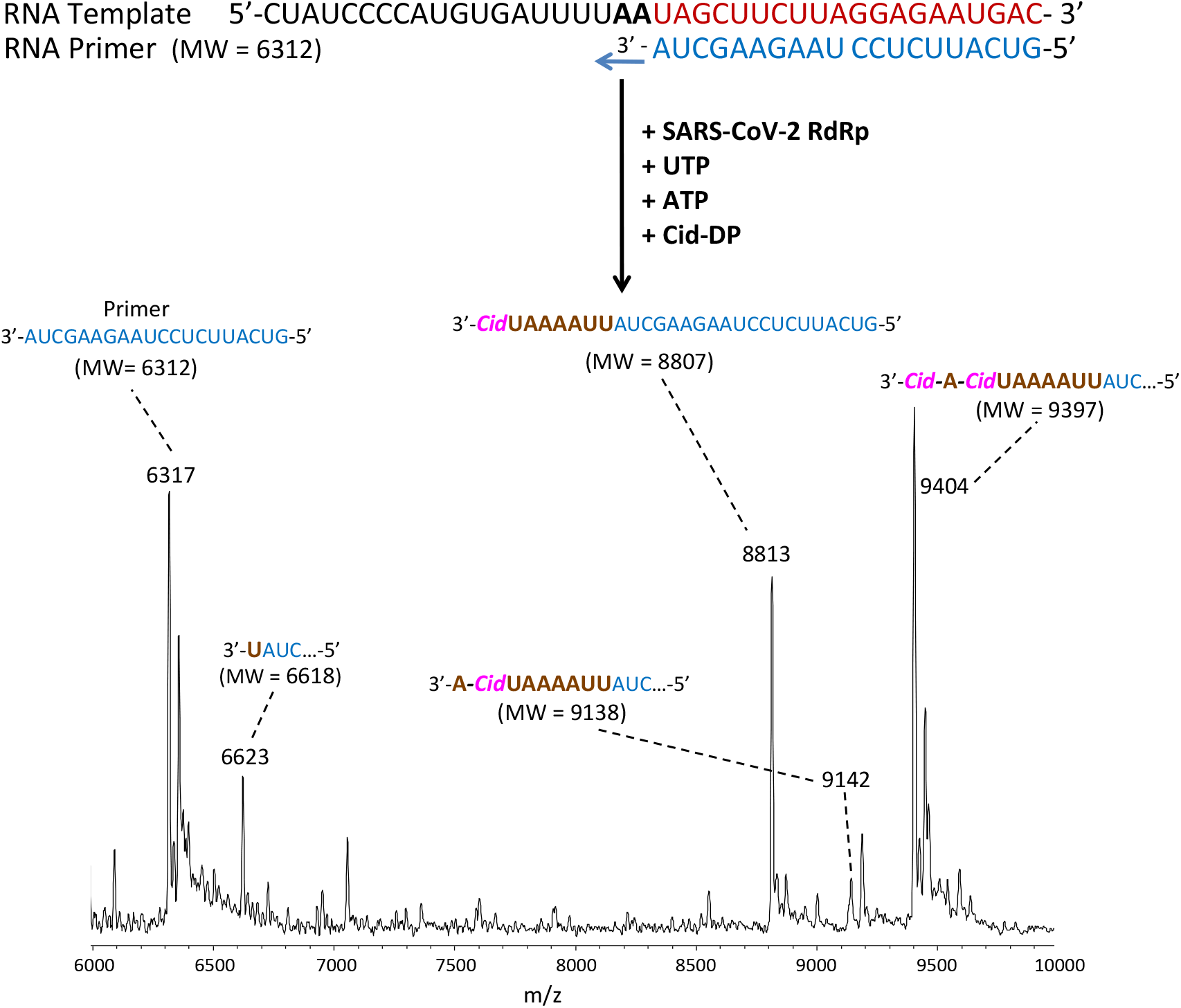
Incorporation of Cid-DP by SARS-CoV-2 RdRp to terminate the polymerase reaction. The sequences of the primer and template used for this extension reaction are shown at the top of the figure. The polymerase extension reaction was performed by incubating Cid-DP, UTP and ATP with preassembled SARS-CoV-2 polymerase (nsp12, nsp7 and nsp8), the indicated RNA template and primer, and the appropriate reaction buffer, followed by detection of reaction products by MALDI-TOF MS. The detailed procedure is shown in the Methods section. The accuracy for m/z determination is ± 10 Da.

The results for the GTP analogues, Car-TP, Ent-TP and Gan-TP are presented in Fig. 5. In each case, extension to the first C position on the template occurs and further extension is blocked in the presence of ATP, UTP and CTP. In more detail, for Car-TP, the major peak observed indicates extension by UTP, ATP and CTP followed by complete termination with a Car-TP (10436 Da observed, 10430 Da expected). In addition, partial extension peaks were seen indicating a single UTP incorporation (6621 Da observed, 6618 Da expected) and extension up to but not including the Car-TP (10123 Da observed, 10121 Da expected). For Ent-TP, a peak was observed indicating extension by UTP, ATP and CTP followed by complete termination by a single Ent-TP (10458 Da observed, 10460 Da expected). Additional peaks are seen representing a single UTP extension (6628 Da observed, 6618 Da expected), and a major peak indicating extension up to but not including the Ent-TP (10129 Da observed, 10121 Da expected). And for Gan-TP, a major peak observed indicated extension by UTP, ATP and CTP followed by complete termination with Gan-TP (10441 Da observed, 10438 Da expected). A small peak representing extension up to but not including Gan-TP (10123 Da observed, 10121 Da expected) was also seen. Similar results were obtained for Car-TP and Gan-TP using the SARS-CoV polymerase (Fig. S3). Both Car-TP and Ent-TP are carbocyclic nucleotides. Car-TP lacks the 2’- and 3’-OH groups, while Ent-TP lacks the 2’-OH group. Gan-TP is an acyclic nucleotide having an OH group but lacking a ribose ring. All three thus may resist the viral exonuclease activity. These results also indicate that Car-TP and Gan-TP are better terminators than Ent-TP; their prodrugs can be evaluated as therapeutics for COVID-19 and SARS.

**Fig. 5.**
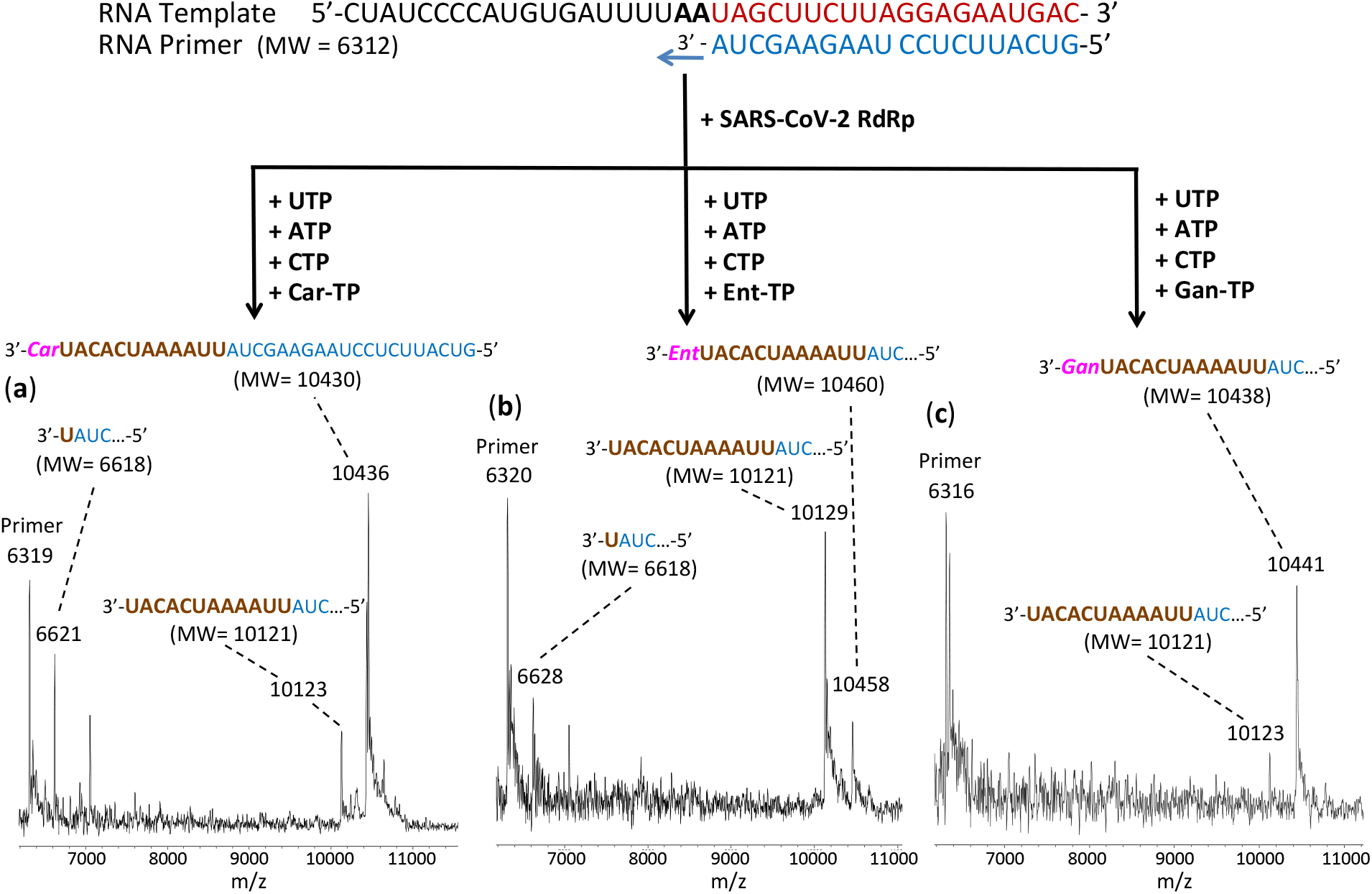
Incorporation of Car-TP, Ent-TP and Gan-TP by SARS-CoV-2 RdRp to terminate the polymerase reaction. The sequences of the primer and template used for this extension reaction are shown at the top of the figure. Polymerase extension reactions were performed by incubating Car-TP, UTP, ATP and CTP (a), Ent-TP, UTP, ATP and CTP (b), and Gan-TP, UTP, ATP and CTP (c) with preassembled SARS-CoV-2 polymerase (nsp12, nsp7 and nsp8), the indicated RNA template and primer, and the appropriate reaction buffer, followed by detection of reaction products by MALDI-TOF MS. The detailed procedure is shown in the Methods section. The accuracy for m/z determination is ± 10 Da.

Fig. S4 shows a side-by-side comparison of the results with 2’-OMe-UTP and 3’-OMe-UTP using the SARS-CoV polymerase. The results for 2’-OMe-UTP are practically identical to those with SARS-CoV-2 in Fig. 4a, indicating that 2’-OMe-UTP exhibits significant polymerase reaction termination. The 3’-OMe-UTP results are consistent with its being an obligate terminator, but with lower incorporation efficiency, represented by a small single-incorporation peak (6625 Da observed, 6632 Da expected).

In Fig. S5, the results are shown for incorporation of 2’-F-dUTP by SARS-CoV RdRp. 2’-F-dUTP was incorporated very efficiently, but also was incorporated opposite the Us in the template strand. This apparent mismatch incorporation may be due to the high concentration of nucleotide analogues used and the relatively low fidelity of SARS-CoV RdRp.

The results for DBiotin-UTP are presented in Fig. S6. DBiotin-UTP incorporation complementary to each A in the template is observed, just like a UTP. Thus, this nucleotide is incorporated and does not terminate the polymerase reaction. These results indicate that modification on the base of the UTP does not affect its incorporation activity by SARS-CoV-2 RdRp.

Fig. S7 presents the results of an experiment where both 2’-OMe-UTP and dUTP were added together at the same concentration. The major peak occurred at 6930 Da (6922 Da expected) representing incorporation by both dUTP and 2’-OMe-UTP in adjacent positions. In addition, some partial extension peaks of a single 2’-OMe-UTP (6626 Da observed, 6632 Da expected) and two dUTPs (6900 Da observed, 6892 Da expected) were found. The incorporation of a dUTP, a 2’-OMe-UTP, both of which lack a 2’-OH group, or their combination would enable them to potentially resist the nsp14 3’-5’ exonuclease activity.

Fig. S8 shows three mass spectra of the polymerase reaction products using equimolar combinations of nucleotide analogues, (a) biotin-dUTP, dUTP, and UTP, (b) 2’-F-dUTP, 2’-OMe-UTP and dUTP, and (c) 2’-NH_2_-dUTP, 2’-OMe-UTP and dUTP, to determine their relative incorporation efficiencies. Based on the results shown in Fig. S8a, biotin-dUTP and dUTP have lower incorporation efficiency than the natural UTP for SARS-CoV-2 RdRp, since peaks are only observed for UTP extension, either one UTP (6620 Da observed, 6618 Da expected) or two UTPs (6928 Da observed, 6924 Da expected). In Fig. S8b, it is seen that 2’-F-dUTP is incorporated far better than 2’-OMe-UTP and dUTP, with the only evident peaks in the spectrum at 6620 Da (6620 Da expected) for extension by one 2’-F-dUTP, and at 6928 Da (6928 Da expected) for extension by two 2’-F-dUTPs. Finally, as shown in Fig. S8c, 2’-NH_2_-dUTP is more efficiently incorporated than 2’-OMe-UTP and dUTP as revealed by the presence of evident peaks only at 6623 Da (6617 Da expected) for extension by one 2’-NH_2_-dUTP and at 6929 Da (6922 Da expected) for extension by two 2’-NH_2_-dUTPs. The results indicate that 2’-F-dUTP and 2’-NH_2_-dUTP behave like UTP, and are not terminators of the polymerase reaction. Neither 2’-F-dUTP nor 2’-NH_2_-dUTP have a free 2’-OH group. It remains to be seen whether the RNAs produced by these two nucleotide analogues will resist exonuclease activity. If that is the case, they may be considered for use in combination with nucleotide analogues that are efficient terminators but sensitive to exonuclease for COVID-19 therapeutic purposes.

In summary, these results demonstrate that the library of nucleotide analogues we tested could be incorporated by the RdRps of SARS-CoV and SARS-CoV-2. Six of these nucleotide analogues exhibited complete termination of the polymerase reaction (3’-OMe-UTP, Car-TP, Gan-TP, Sta-TP, Ent-TP, Biotin-dUTP), two showed incomplete or delayed termination (2’-OMe-UTP, Cid-DP), and 3 did not terminate the polymerase reaction (2’-F-dUTP, 2’-NH_2_-dUTP and DBiotin-UTP) using the RdRp of SARS-CoV and/or SARS-CoV-2. Their prodrug versions (shown in Fig. 2 and S1) are available or can be readily synthesized using the ProTide approach (Alanazi et al. 2019). The prodrugs of several of these nucleoside triphosphate analogues (Ganciclovir/Valganciclovir, Cidofovir/Brincidofovir, Abacavir, Stavudine, and Entecavir) have been FDA approved for treatment of other viral infections and their toxicity profile is well established. Thus, our results provide a molecular basis for further evaluation of these prodrugs in SARS-CoV-2 virus inhibition and animal models to test their efficacy for the development of potential COVID-19 therapeutics. To reduce drug resistance, it would be useful to test them in combination protocols.

## Author contributions

J.J. and R.N.K. conceived and directed the project; the approaches and assays were designed and conducted by J.J., X.L., S.K., S.J., J.J.R., M.C. and C.T., and SARS-CoV/SARS-CoV-2 polymerases (nsp12) and associated proteins (nsp 7 and 8) were cloned and purified by T.K.A. and R.N.K. Data were analyzed by all authors. All authors wrote and reviewed the manuscript.

## Supplementary Material

**Fig. S1.**
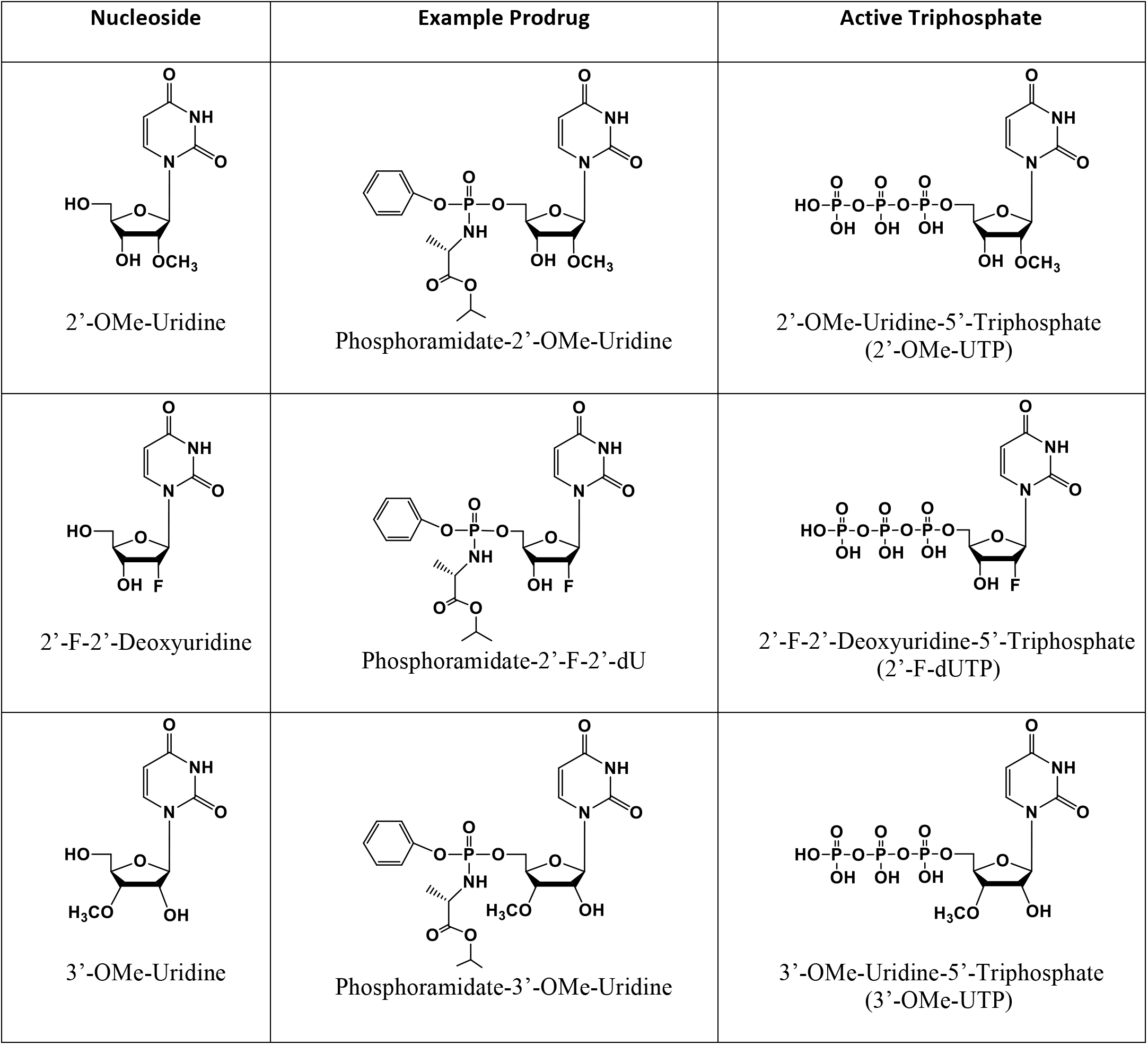
Structures of nucleoside analogues, example prodrugs and active triphosphate forms. The nucleosides 2’-OMe-Uridine, 2’-F-2’-Deoxyuridine and 3’-OMe-Uridine (left), example prodrug forms (middle) and their active triphosphate forms (right).

**Fig. S2.**
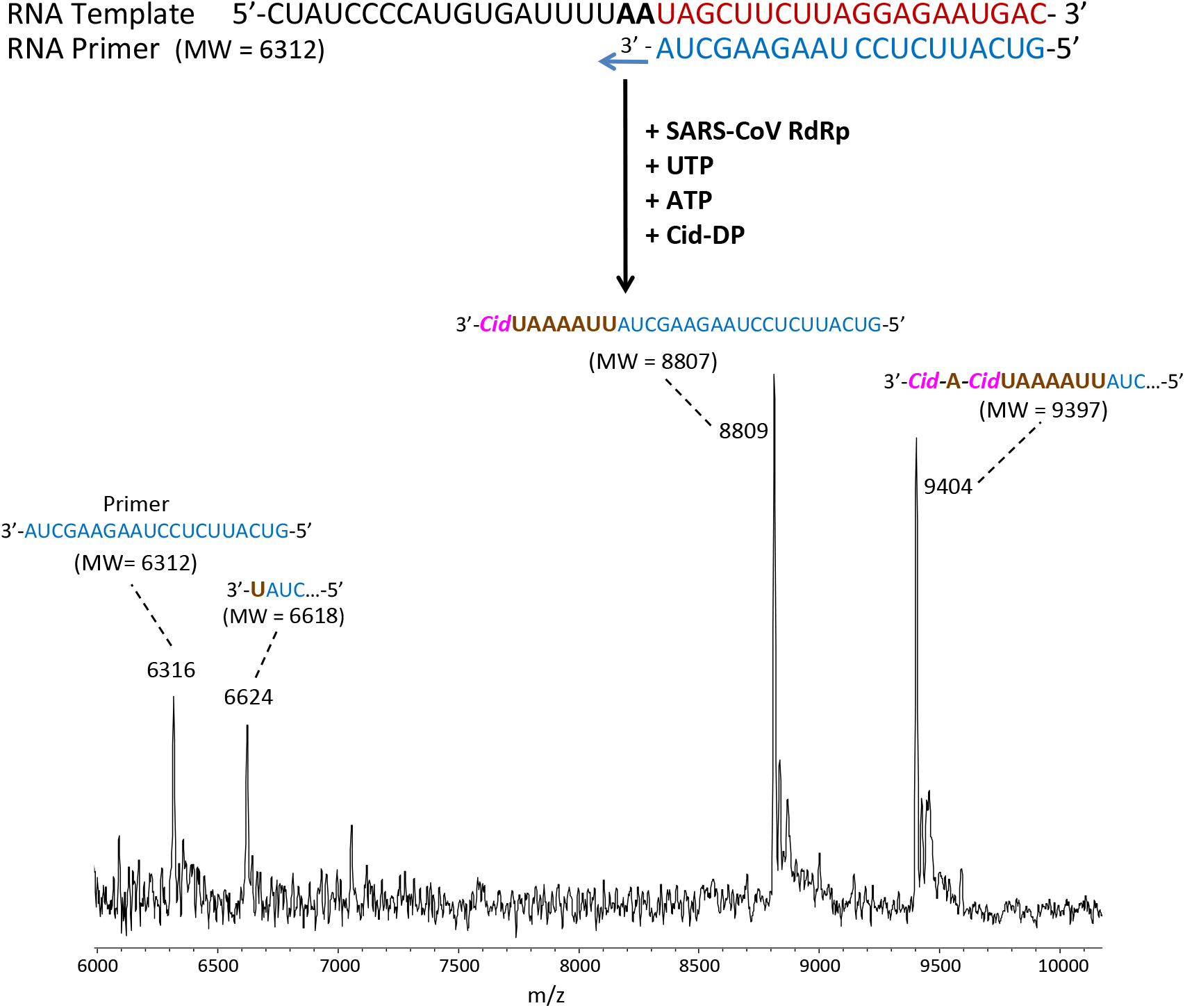
Incorporation of Cid-DP by SARS-CoV RdRp to terminate the polymerase reaction. The sequences of the primer and template used for this extension reaction are shown at the top of the figure. The polymerase extension reaction was performed by incubating Cid-DP, UTP and ATP with preassembled SARS-CoV polymerase (nsp12, nsp7 and nsp8), the indicated RNA template and primer, and the appropriate reaction buffer, followed by detection of reaction products by MALDI-TOF MS. The detailed procedure is shown in the Methods section. The accuracy for m/z determination is ± 10 Da.

**Fig. S3.**
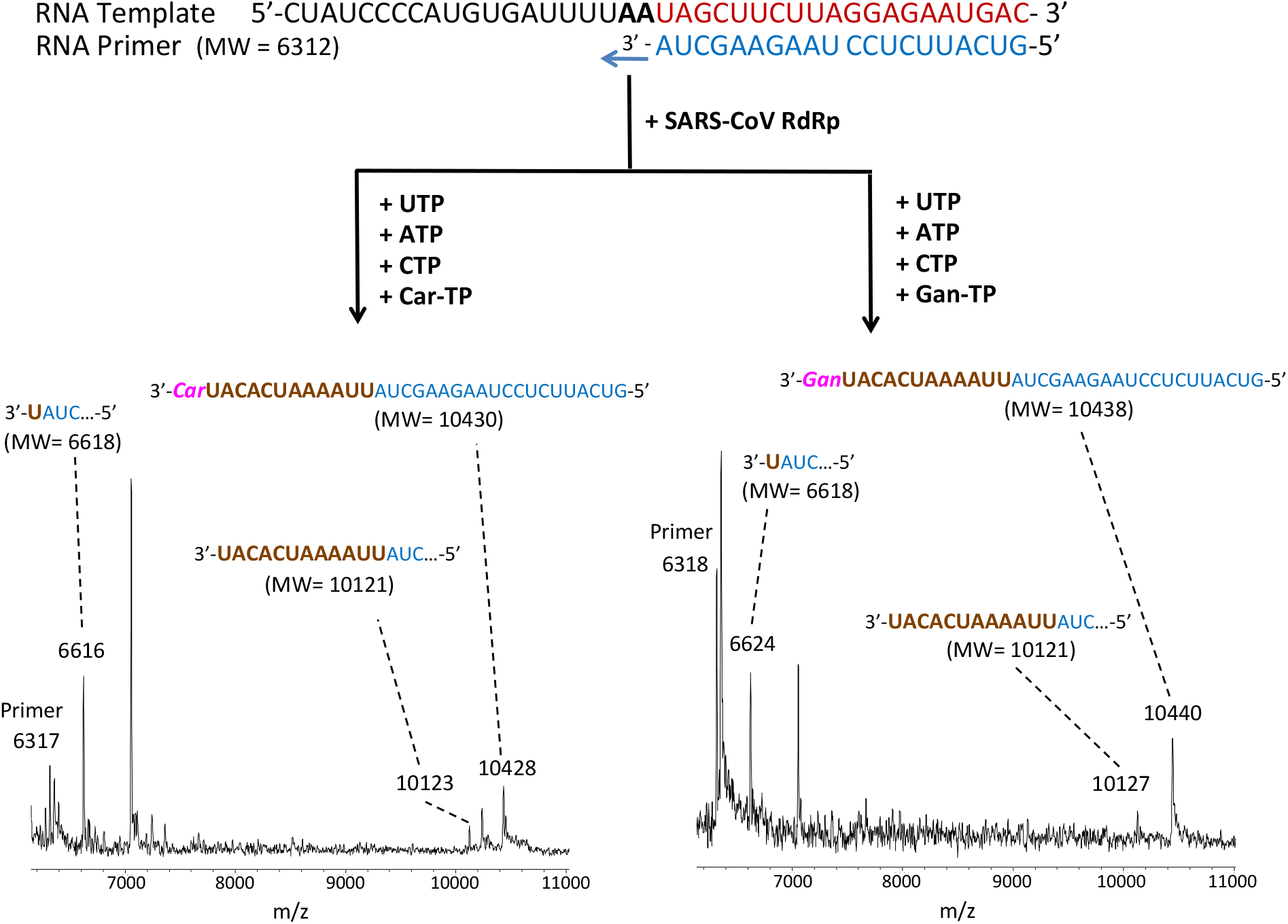
Incorporation of Car-TP and Gan-TP by SARS-CoV RdRp to terminate the polymerase reaction. The sequences of the primer and template used for this extension reaction are shown at the top of the figure. Polymerase extension reactions were performed by incubating Car-TP, UTP, ATP and CTP (left) and Gan-TP, UTP, ATP and CTP (right) with pre-assembled SARS-CoV polymerase (nsp12, nsp7 and nsp8), the indicated RNA template and primer, and the appropriate reaction buffer, followed by detection of reaction products by MALDI-TOF MS. The detailed procedure is shown in the Methods section. The accuracy for m/z determination is ± 10 Da.

**Fig. S4.**
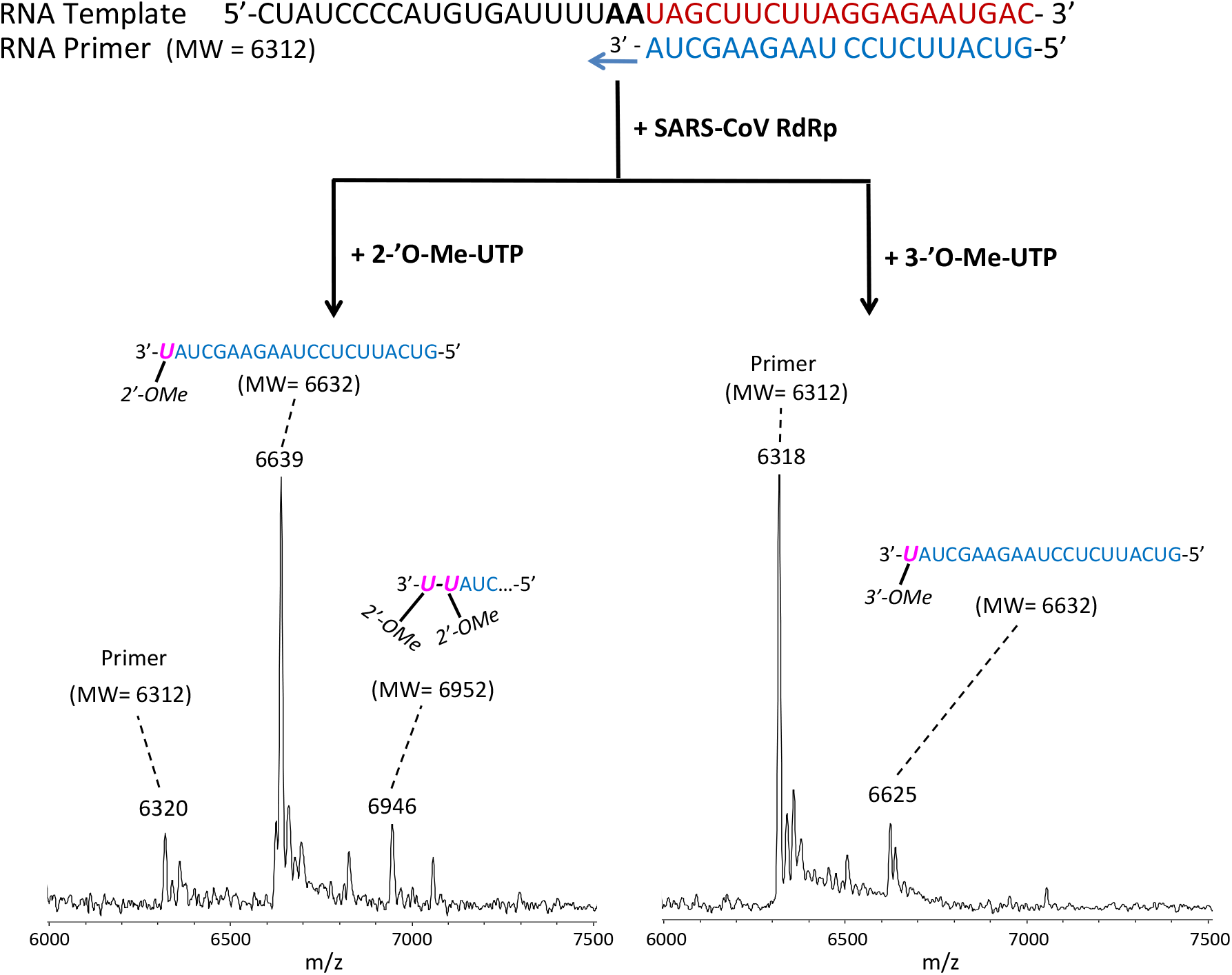
Incorporation of 2’-OMe-UTP and 3’-OMe-UTP by SARS-CoV RdRp to terminate the polymerase reaction. The sequences of the primer and template used for this extension reaction are shown at the top of the figure. Polymerase extension reactions were performed by incubating 2’-OMe-UTP (left) and 3’-OMe-UTP (right) with pre-assembled SARS-CoV polymerase (nsp12, nsp7 and nsp8), the indicated RNA template and primer, and the appropriate reaction buffer, followed by detection of reaction products by MALDI-TOF MS. The detailed procedure is shown in the Methods section. The accuracy for m/z determination is ± 10 Da.

**Fig. S5.**
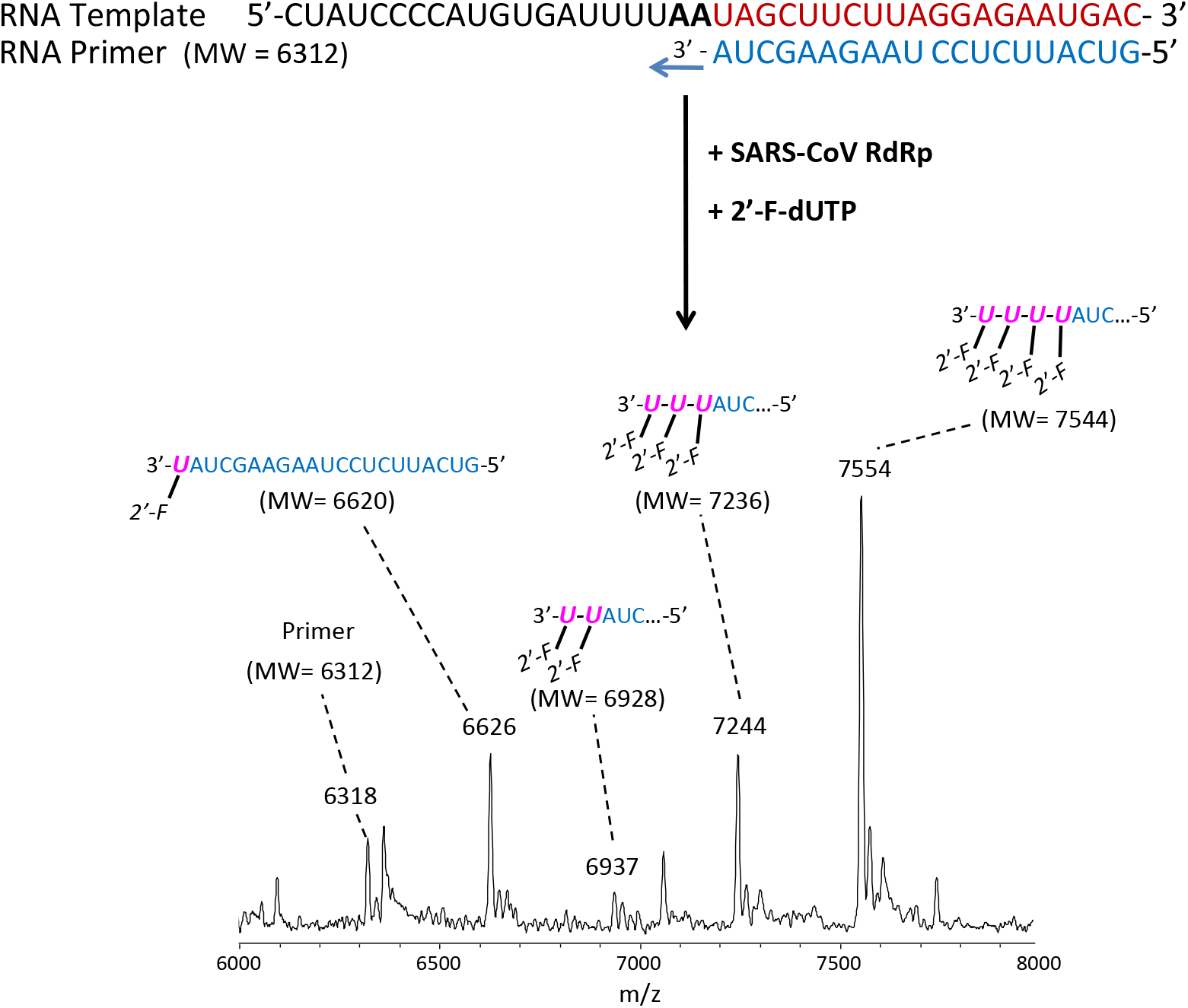
Incorporation of 2’-F-dUTP by SARS-CoV RdRp catalyzed reaction. The sequences of the primer and template used for this extension reaction are shown at the top of the figure. The polymerase extension reaction was performed by incubating 2’-F-dUTP with pre-assembled SARS-CoV polymerase (nsp12, nsp7 and nsp8), the indicated RNA template and primer, and the appropriate reaction buffer, followed by detection of reaction products by MALDI-TOF MS. The detailed procedure is shown in the Methods section. The accuracy for m/z determination is ± 10 Da.

**Fig. S6.**
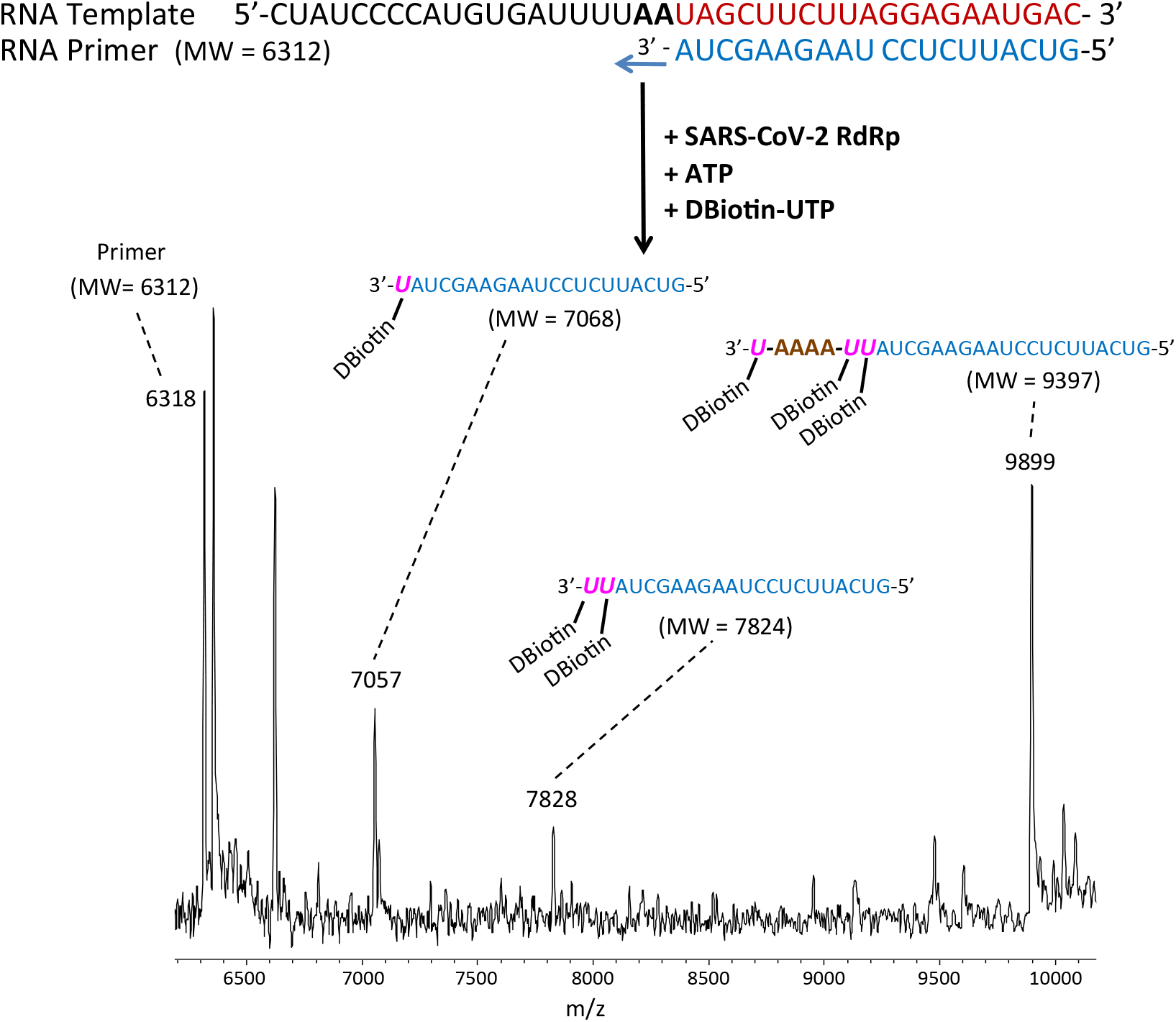
Incorporation of DBiotin-UTP by SARS-CoV-2 RdRp catalyzed reaction. The sequences of the primer and template used for this extension reaction are shown at the top of the figure. The polymerase extension reaction was performed by incubating DBiotin-UTP and ATP with pre-assembled SARS-CoV-2 polymerase (nsp12, nsp7 and nsp8), the indicated RNA template and primer, and the appropriate reaction buffer, followed by detection of reaction products by MALDI-TOF MS. The detailed procedure is shown in the Methods section. The accuracy for m/z determination is ± 10 Da.

**Fig. S7.**
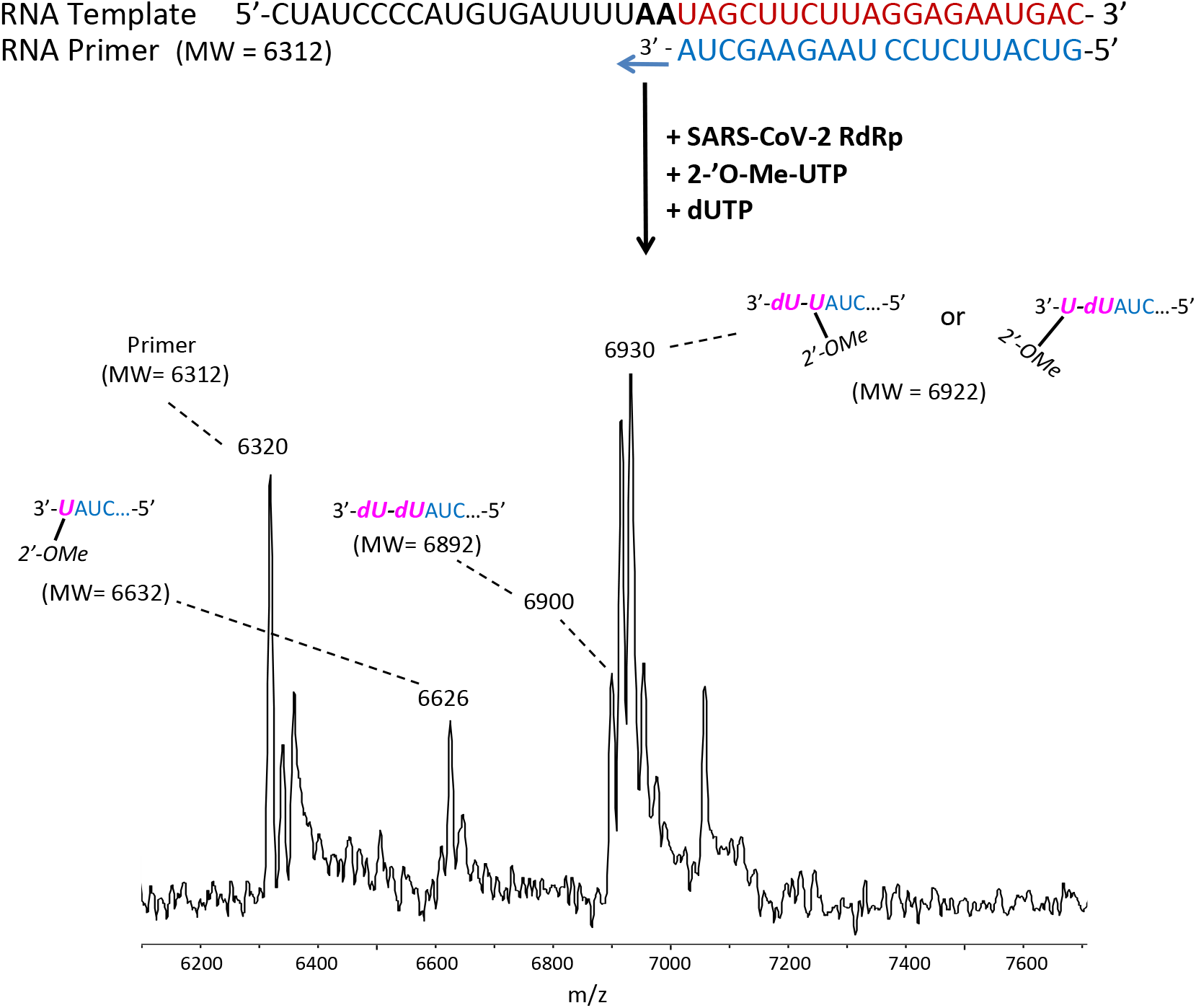
Incorporation of 2’-OMe-UTP and dUTP by SARS-CoV-2 RdRp catalyzed reaction. The sequences of the primer and template used for this extension reaction are shown at the top of the figure. The polymerase extension reaction was performed by incubating 2’-OMe-UTP and dUTP with preassembled SARS-CoV-2 polymerase (nsp12, nsp7 and nsp8), the indicated RNA template and primer, and the appropriate reaction buffer, followed by detection of reaction products by MALDI-TOF MS. The detailed procedure is shown in the Methods section. The accuracy for m/z determination is ± 10 Da.

**Fig. S8.**
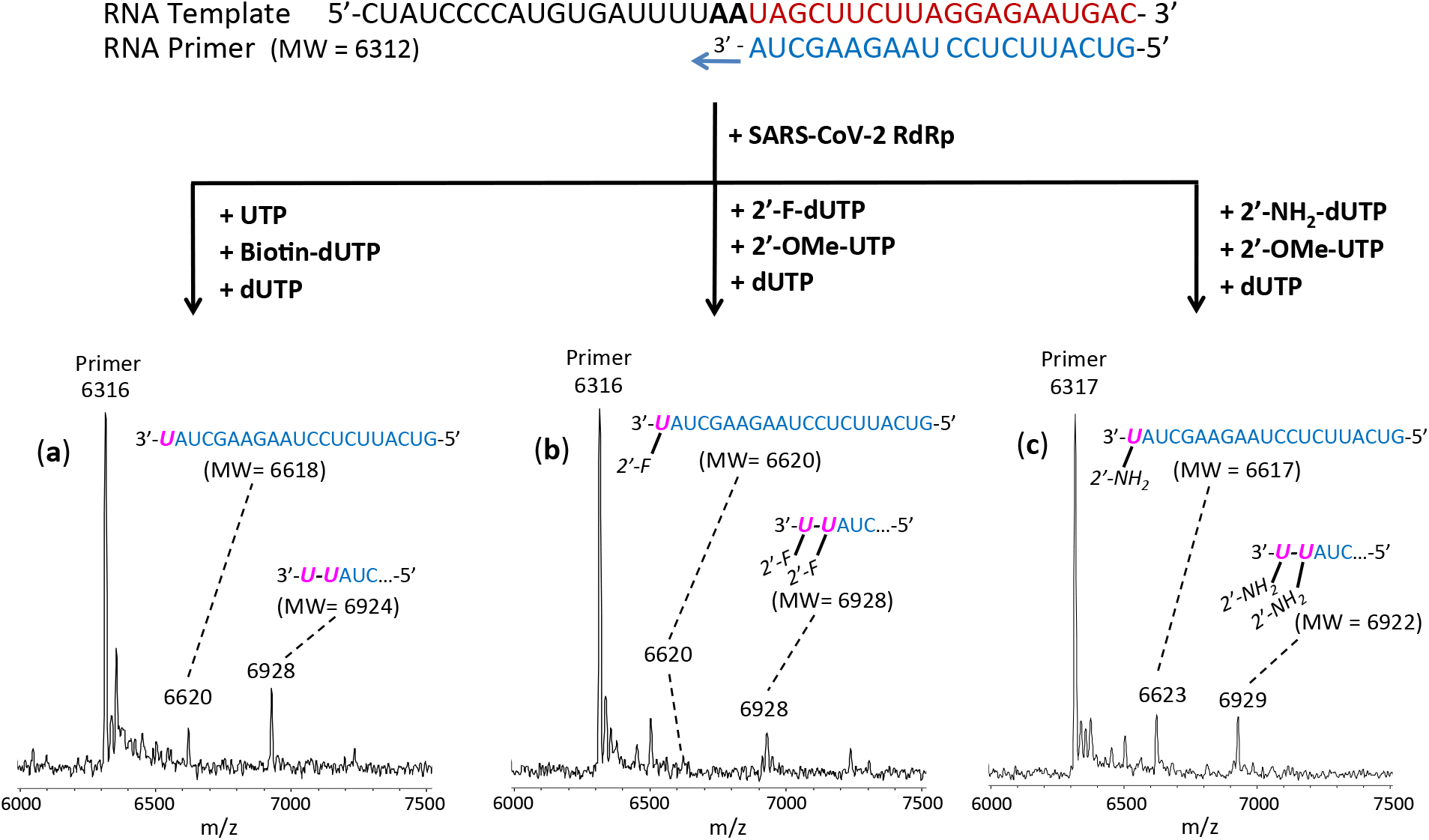
Comparison of Relative Incorporation of UTP, Biotin-dUTP, dUTP; 2’-F-dUTP, 2’-OMe-UTP, dUTP; and 2’-NH_2_-dUTP, 2’-OMe-UTP, dUTP by SARS-CoV-2 RdRp. The sequences of the primer and template used for this extension reaction are shown at the top of the figure. Polymerase extension reactions were performed by incubating UTP, Biotin-dUTP and dUTP (a), 2’-F-dUTP, 2’-OMe-UTP and dUTP (b) and 2’-NH2-dUTP, 2’-OMe-UTP and dUTP (c) with pre-assembled SARS-CoV-2 polymerase (nsp12, nsp7 and nsp8), the indicated RNA template and primer, and the appropriate reaction buffer, followed by detection of reaction products by MALDI-TOF MS. The detailed procedure is shown in the Methods section. The accuracy for m/z determination is ± 10 Da.

## References

Agostini, M.L. et al., 2019. Small-molecule antiviral β-D-N^4^-hydroxycytidine inhibits a proofreading-intact coronavirus with a high genetic barrier to resistance. J. Virol. 93, e01348–e01319. https://doi.org/10.1128/JVI.01348-19.

Akyürek, L.M. et al., 2001. Coexpression of guanylate kinase with thymidine kinase enhances prodrug cell killing *in vitro* and suppresses vascular smooth muscle cell proliferation *in vivo*. Molec. Ther. 3, 779–786. https://doi.org/10.1006/mthe.2001.0315.

Alanazi, A.S., James, E., Mehellou, Y., 2019. The ProTide prodrug technology: where next? ACS Med. Chem. Lett. 10, 2–5. https://doi.org/10.1021/acsmedchemlett.8b00586.

Anderson, P.L., Kiser, J.J., Gardner, E.M., Rower, J.E., Meditz, A., Grant, R.M., 2011. Pharmacological considerations for tenofovir and emtricitabine to prevent HIV infection. J. Antimicrob. Chemother. 66, 240–250. https://doi.org/10.1093/jac/dkq447.

Arup, H., Williams, D.M., Eckstein, F., 1992. 2’-Fluoro and 2,-amino-2’-deoxynucleoside 5’-triphosphates as substrates for T7 RNA polymerase. Biochem. 31, 9636–9641. https://doi.org/10.1021/bi00155a016.

Bouvet, M., Imbert, I., Subissi, L., Gluais, L., Canard, B., Decroly, E., 2012. RNA 3’-end mismatch excision by the severe acute respiratory syndrome coronavirus nonstructural protein nsp10/nsp14 exoribonuclease complex. Proc. Natl. Acad. Sci. USA 109, 9372–9377. https://doi.org/10.1073/pnas.1201130109.

Chien, M. et al., 2020. Nucleotide analogues as inhibitors of SARS-CoV-2 polymerase. bioRxiv. https://doi.org/10.1101/2020.03.18.997585.

Cundy, K.C., 1999. Clinical pharmacokinetics of the antiviral nucleotide analogues cidofovir and adefovir. Clin. Pharmacokinet. 36, 127–143. https://doi.org/10.2165/00003088-199936020-00004.

Dustin, L.B., Bartolini, B., Capobianchi, M.R., Pistello, M., 2016. Hepatitis C virus: life cycle in cells, infection and host response, and analysis of molecular markers influencing the outcome of infection and response to therapy. Clin. Microbiol. Infect. 22, 826–832. https://doi.org/10.1016/j.cmi.2016.08.025.

De Clercq, E., 2002. Cidofovir in the treatment of poxvirus infections. Antivir. Res. 55, 1–13. https://doi.org/10.1016/S0166-3542(02)00008-6.

De Clercq, E., Li, G., 2016. Approved antiviral drugs over the past 50 years. Clin. Microbiol. Rev. 29, 695–747. https://doi.org/10.1128/CMR.00102-15.

Eckerle, L.D. et al., 2010. Infidelity of SARS-CoV nsp14-exonuclease mutant virus replication is revealed by complete genome sequencing. PLoS Pathogens 6, e1000896. https://doi.org/10.1371/journal.ppat.1000896.

Elfiky, A.A., 2020, Ribavirin, Remdesivir, Sofosbuvir, Galidesivir, and Tenofovir against SARS-CoV-2 RNA dependent RNA polymerase (RdRp): A molecular docking study. Life Sciences. 253, 117592. https://doi.org/10.1016/j.lfs.2020.117592.

Eltahla, A.A., Luciani, F., White, P.A., Lloyd, A.R., Bull, R.A., 2015. Inhibitors of the hepatitis C virus polymerase; mode of action and resistance. Viruses 7, 5206–5224. https://doi.org/10.3390/v7102868.

Eyer, L., Fojtíková, M., Nencka, R., Rudolf, I., Hubálek, Z., Růzěk, D., 2019. Viral RNA-dependent RNA polymerase inhibitor 7-deaza-2’-C-methyladenosine prevents death in a mouse model of West Nile virus infection. Antimicrob. Agents Chemother. 63, e02093–18. https://doi.org/10.1128/AAC.02093-18.

Faletto, M.B., Miller, W.H., Garvey, E.P., St. Clair, M.H., Daluge, S.M., Good, S.S., 1997. Unique intracellular activation of the potent anti-human immunodeficiency virus agent 1592U89. Antimicrob. Agents Chemother. 41, 1099–1107. https://doi.org/10.1128/aac.4L5.1099.

Ferron, F. et al., 2018. Structural and molecular basis of mismatch correction and ribavirin excision from coronavirus RNA. Proc. Natl. Acad. Sci. USA, 115, E162–E171. https://doi.org/10.1073/pnas.1718806115.

Gao, Y. et al., 2020. Structure of the RNA-dependent RNA polymerase from COVID-19 virus. Science. https://doi.org/10.1126/science.abb7498.

Gordon, C.J., Tchesnokov, E.P., Feng, J.Y., Porter, D.P., Götte, M., 2020. The antiviral compound remdesivir potently inhibits RNA-dependent RNA polymerase from Middle East respiratory syndrome coronavirus. J. Biol. Chem. 295, 4773–4779. https://doi.org/10.1074/jbc.AC120.013056.

Gordon, C.J. et al., 2020. Remdesivir is a direct-acting antiviral that inhibits RNA-dependent RNA polymerase from severe acute respiratory syndrome coronavirus 2 with high potency. J. Biol. Chem. https://doi.org/10.1074/jbc.RA120.013679.

Ho, H-T., Hitchcock, M.J.M., 1989. Cellular pharmacology of 2’,3’-dideoxy-2’,3’-didehydrothymidine, a nucleoside analog active against human immunodeficiency virus. Antimicrob. Agents Chemother. 33, 844–849. https://doi.org/10.1128/AAC.33.6.844.

Huang, P., Farquhar, D., Plunkett, W., 1992. Selective action of 2’,3’-didehydro-2’,3’-dideoxythymidine triphosphate on human immunodeficiency virus reverse transcriptase and human DNA polymerases. J. Biol. Chem. 267, 2817–2822. https://www.jbc.org/content/267/4/2817.full.pdf.

Jockusch, S. et al., 2020. Triphosphates of the two components in DESCOVY and TRUVADA are inhibitors of the SARS-CoV-2 polymerase. bioRxiv. https://doi.org/10.1101/2020.04.03.022939.

Ju, J. et al., 2020. Nucleotide analogues as inhibitors of SARS-CoV polymerase. bioRxiv. https://doi.org/10.1101/2020.03.12.989186.

Kirchdoerfer, R.N., Ward, A.B., 2019. Structure of the SARS-CoV nsp12 polymerase bound to nsp7 and nsp8 co-factors. Nature Commun. 10, 2342. https://doi.org/10.1038/s41467-019-10280-3.

Kumaki, Y. et al., 2011. *In vitro* and *in vivo* efficacy of fluorodeoxycytidine analogs against highly pathogenic avian influenza H5N1, seasonal, and pandemic H1N1 virus infections. Antiviral Res. 9, 329–340. https://doi.org/10.1016/j.antiviral.2011.09.001.

Lanier, R. et al., 2010. Development of CMX001 for the treatment of poxvirus infections. Viruses 2, 2740–2762. https://doi.org/10.3390/v2122740.

Lauridsen, L.H., Rothnagel, J.A., Veedu, R.N., 2012. Enzymatic recognition of 2’-modified ribonucleoside 5’-triphosphates: Towards the evolution of versatile aptamers. ChemBioChem. 13, 19–25. https://doi.org/10.1002/cbic.201100648.

Ma, Y. et al., 2015. Structural basis and functional analysis of the SARS coronavirus nsp14-nsp10 complex. Proc. Natl. Acad. Sci. USA. 112, 9436–9441. https://doi.org/10.1073/pnas.1508686112.

Magee, W.C., Hostetler, K.Y., Evans, D.H., 2005. Mechanism of inhibition of vaccinia virus DNA polymerase by cidofovir diphosphate. Antimicrob. Agents Chemother. 49, 3153–3162. https://doi.org/10.1128/AAC.49.8.3153-3162.2005.

Matthews, S.J., 2006. Entecavir for the treatment of chronic hepatitis B virus infection. Clin. Therapeut. 28, 184–203. https://doi.org/10.1016/j.clinthera.2006.02.012.

Matthews, T., Boehme, R., 1988. Antiviral activity and mechanism of action of ganciclovir. Rev. Infect. Dis. 10, S490–S494. https://doi.org/10.1093/clinids/10.supplement_3.s490.

Mazzucco, C.E., Hamatake, R.K., Colonno, R.J., Tenney, D.J., 2008. Entecavir for treatment of hepatitis B virus displays no in vitro mitochondrial toxicity or DNA polymerase gamma inhibition. Antimicrob. Agents Chemother. 52, 598–605. https://doi.org/10.1128/AAC.01122-07.

McKenna, C.E. et al., 1989. Inhibitors of viral nucleic acid polymerases. Pyrophosphate analogues. ACS Symposium Series. 401, 1–16. Chapter 1. https://doi.org/10.1021/bk-1989-0401.ch001.

Minskaia, E. et al., 2006. Discovery of an RNA virus 3’-->5’ exoribonuclease that is critically involved in coronavirus RNA synthesis. Proc. Natl. Acad. Sci. USA. 103, 5108–5113. https://doi.org/10.1073/pnas.0508200103.

Öberg, B. (2006) Rational design of polymerase inhibitors as antiviral drugs. Antiviral Res. 71, 90–95. https://doi.org/10.1016/j.antiviral.2006.05.012.

Ray, A.S., Basavapathruni, A., Anderson, K.S., 2002. Mechanistic studies to understand the progressive development of resistance in human immunodeficiency virus type 1 reverse transcriptase to abacavir. J. Biol. Chem. 277, 40479–40490. https://doi.org/10.1074/jbc.M205303200.

Rivkina, A., Rybalov, S., 2002. Chronic hepatitis B: current and future treatment options. Pharmacother. 22, 721–737. https://doi.org/10.1592/phco.22.9.721.34058.

Selisko, B., Papageorgiou, N., Ferron, F., Canard, B., 2018. Structural and functional basis of the fidelity of nucleotide selection by *Flavivirus* RNA-dependent RNA polymerases. Viruses. 10, 59. https://doi.org/10.3390/v10020059.

Shannon, A. et al., 2020. Remdesivir and SARS-CoV-2: Structural requirements at both nsp12 RdRp and nsp14 exonuclease active-sites. Antivir Res 178, 104793. https://doi.org/10.1016/j.antiviral.2020.104793.

Sheahan, T.P. et al., 2020. An orally bioavailable broad-spectrum antiviral inhibits SARS-CoV-2 in human airway epithelial cell cultures and multiple coronaviruses in mice. Sci. Transl. Med. https://doi.org/10.1126/scitranslmed.abb5883.

Subissi, L. et al., 2014. One severe acute respiratory syndrome coronavirus protein complex integrates processive RNA polymerase and exonuclease activities. Proc. Natl. Acad. Sci. USA. 111, E3900–E3909. https://doi.org/10.1073/pnas.1323705111.

Tchesnokov, E.P. et al., 2008. Delayed chain termination protects the anti-hepatitis B virus drug entecavir from excision by HIV-1 reverse transcriptase. J. Biol. Chem. 283, 34218–34228. https://doi.org/10.1074/jbc.M806797200.

te Velthuis, A.J.W., 2014. Common and unique features of viral RNA-dependent polymerases. Cell Mol. Life Sci. 71, 4403–4420. https://doi.org/10.1007/s00018-014-1695-z.

Trost, L.C. et al., 2015. The efficacy and pharmacokinetics of brincidofovir for the treatment of lethal rabbitpox virus infection: A model of smallpox disease. Antivir. Res. 117, 115–121. https://doi.org/10.1016/j.antiviral.2015.02.007.

Zhu, N. et al., 2020. A novel coronavirus from patients with pneumonia in China, 2019. N. Eng. J. Med. 382, 727–733. https://doi.org/10.1056/NEJMoa2001017.

Zumla, A., Chan, J.F.W., Azhar, E.I., Hui, D.S.C., Yuen, K.-Y. 2016. Coronaviruses - drug discovery and therapeutic options. Nat. Rev. Drug Discovery. 15, 327–347. https://doi.org/10.1038/nrd.2015.37.

